# Learning-Augmented Sketching Offers Improved Performance for Privacy Preserving and Secure GWAS

**DOI:** 10.1101/2024.09.19.613975

**Authors:** Junyan Xu, Kaiyuan Zhu, Jieling Cai, Can Kockan, Natnatee Dokmai, Hyunghoon Cho, David P. Woodruff, S. Cenk Sahinalp

## Abstract

The introduction of trusted execution environments (TEEs), such as secure enclaves provided by the Intel SGX technology has enabled secure and privacy-preserving computation on the cloud. The stringent resource limitations, such as memory constraints, required by some TEEs necessitates the development of computational approaches with reduced memory usage, such as sketching. One example is the SkSES method for GWAS on a cohort of case and control samples from multiple institutions, which identifies the most significant SNPs in a privacy-preserving manner without disclosing sensitive genotype information to other institutions or the cloud service provider. Here we show how to improve the performance of SkSES on large datasets by augmenting it with a learning-augmented approach. Specifically, we show how individual institutions can perform smaller scale GWAS on their own datasets and identify two sets of variants according to certain criteria, which are then used to guide the sketching process to more accurately identify significant variants over the collective dataset. The new method achieves up to 40% accuracy gain compared to the original SkSES method under the same memory constraints on datasets we tested on. The code is available at https://github.com/alreadydone/sgx-genome-variants-search.

## 1 Introduction

The ongoing expansion and increasing significance of genomic datasets necessitate the development of privacy-preserving and secure methods to process and analyze them. Genomic data carries sensitive insights into an individual’s predisposition to diseases [1], extending potential privacy implications to their biological relatives [2]. The inherent immutability of genomic information and the potential for unforeseen privacy breaches in the future [3] highlight the criticality of safeguarding this data. As a result, policies or law could severely limit or fully prohibit exchange of genomic data between hospitals, research institutions, government organizations and countries. A number of cryptographic techniques [4, 5, 6, 7, 8, 9, 10, 11] have been proposed to address privacy concerns related to genomic data, but their high computational overhead means they cannot scale up to offer the performance needed for real-life genomic analysis [12, 13, 14, 15].

A genome-wide association study (GWAS) aims to identify genetic variants associated with a particular trait (e.g. disease), and it does so by computing a certain statistic for each variant over a large cohort of cases and controls. Bringing together such a cohort, especially for rare diseases, may necessitate multiple institutions, possibly from several countries, to share data and collectively develop an accurate model. SkSES (sketching algorithms for secure-enclave-based genomic data analysis) [16] is a computational framework to perform secure collaborative GWAS in an untrusted cloud platform with minimal performance overhead, through the use of Intel’s Software Guard Extensions (SGX) [17], a combined hardware and software platform supported by all current-generation Intel processors, that offers users sensitive data analysis within a protected enclave. Each sample’s genotype vector is encrypted and uploaded independently to the secure enclave, and it is decrypted there, so no institution nor the cloud provider could gain access to the sensitive genomic data, alleviating privacy concerns for such large collaborative genomic analysis.

Unlike software-based techniques such as the garbled-circuit-based FlexSC framework [18] or the homomorphic encryption-based HElib framework [19], the SGX platform does not introduce substantial computational overhead or restrictions on basic operations, and thus is likely to make secure genome-scale data analysis feasible. However, it does come with memory limitations, which SkSES addresses by a “sketching” approach to summarize variant frequencies within the limited memory of the secure enclave and identify the most significant variants across case and control samples (i.e., those that best differentiate the case and control samples) according to the *χ*^2^ statistic.

One key challenge in GWAS is sorting out systematic differences between different human subpopulations [20]: disease-causing genetic changes may get passed down alongside harmless ones, leading to false hits. This confounding effect is known as population stratification and requires mitigation using methods such as EIGENSTRAT [21], which subtracts off principal components identified through PCA. Due to limited memory of the enclave, sketching is also applied by SkSES to perform PCA.

As the size of the dataset increases, the sketches become more crowded, and the estimation error soars. We hypothesize that “collisions”, i.e., multiple significant (in terms of the statistic used by the GWAS or another quantity used as a proxy for the statistic) SNPs being assigned to the same entry (i.e., “bucket”) in the sketches, might be a main contributing factor to the error. In this paper, we address this issue by introducing a learning-augmented approach to identify the most significant SNPs in a training dataset and assign them to “unique buckets” outside of the sketches; this also reduces their impact on the estimated quantity for SNPs that are non-significant. In addition, we show how to identify another set of representative SNPs that are most useful for accurately identifying principal components during PCA for reducing the impact of population stratification via EIGENSTRAT. These SNPs are identified by running GWAS on a much smaller dataset (say 10% of the size of the full dataset), which can be done much less costly and locally by each or any of the data-holding parties involved. According to our tests, by employing these two sets of SNPs, our learning-augmented version of SkSES significantly improves the accuracy of the original version of SkSES without introducing any overhead with respect to memory usage or speed.

## 2 Methods

### 2.1 Preliminaries

#### 2.1.1 The SGX platform in a nutshell

Intel SGX (Software Guard Extensions) is a commonly-used Trusted Execution Environment (TEE) technology embedded in modern Intel CPU models, which offers hardware-based protection for applications running inside an SGX *enclave* - an isolated and protected computing environment against adversaries controlling the host operating system. The security model provided by SGX hardware can prevent adversaries from learning private data or modifying the control flow of a protected application without compromising the hardware itself. As such a common use case of SGX is for secure *outsourced* computation, in which a user with limited computational capabilities sends private data to a remote SGX enclave, controlled by an untrusted party, for secure data processing. This enticing possibility has been gathering increasing attention in the bioinformatics community in recent years [22, 23, 24, 25, 26, 27, 16, 28, 29, 30] due to its potential to accelerate scientific discoveries by facilitating the sharing of sensitive biomedical data. For security in the outsourcing scenario, SGX usually employs a cryptographic protocol called *remote attestation* to provide proof to a remote user that both the enclave environment and the application running inside the enclave would not be tampered with; and that the communication channel for the transfer of sensitive data is secure.

In this work, we focus on the **algorithmic improvement** on the GWAS protocol but do not address the potential side-channel leaks which refer to externally measurable behaviors of the program (e.g. runtime or memory access patterns) that may be used to infer the underlying sensitive data. Note that side-channel attacks can be usually addressed with an oblivious implementation of the algorithm but at the cost of increased running time.

#### 2.1.2 An overview of the SkSES method for identifying the top SNPs in an SGX enclave

We focus on the most general GWAS protocol introduced by SkSES [16] involving three steps: (i) Population stratification correction using PCA; (ii) Top *l* SNP candidate identification by sketching; and (iii) Top *k* SNP identification among the *l* candidates. The computation process involves a server equipped with Intel SGX hardware, and the client(s) that want to perform collaborative GWAS in a privacy-preserving and secure manner. Each of the three steps requires the client(s) to encrypt and send the VCF files they store in a compressed format by only keeping the SNP IDs and their zygosity status. The server, i.e., the SGX enclave memory maintains a distinct data structure for each step, which is updated online after the server receives each compressed and encrypted VCF file from the clients. In each step, once all incoming data have been processed, the server postprocess the data structure and computes all necessary intermediate results required for the next step (or the final output). The size of these intermediate results is usually small (negligible compared to the data structures) and thus are stored independently (from the data structures) in the enclave memory. Finally, after the last step the results consisting of the top *k* significant SNPs according to the desired association test are sent back to the clients. We describe each of the steps in more detail below.

i. **Population stratification correction using PCA**. In this step the server constructs a sketch genotype matrix *G′* which corresponds to a random projection (of the columns) of the underlying genotype matrix *G* given by the input individuals (VCF files). Specifically, *G* ∈ {0, 1, 2}^*m×n*^ denotes the genotype matrix where *m* is the number of VCF files and *n* is the total number of SNPs included in the *m* samples; *G′* = *GK* denotes the sketch matrix of size *m × n′* (*n′ ≪ n*) which can fit in the enclave memory, where *K* is a sparse count-sketch matrix of size *n × n′* with exactly one random *±*1 entry in each row. A compressed and encrypted VCF file encodes all non-zero entries in a particular row **g**_*j*_ ∈ {0, 1, 2}^*n*^ in *G*. The update of *G′* from the *j*-th individual simply involves adding a vector **g**_*j*_ · *K* to the *j*-th row of *G′*. After all encrypted VCF files are processed, the server first normalize each column of *G′* by subtracting the mean and optionally normalizing by the standard deviation. Let the normalized matrix be *X′*. Then SVD is performed on *X′* to compute the top *y* left singular vectors *U*_*y*_ of size *m × y*, which will be stored in the enclave memory and used in the following steps of computation. In addition, the phenotype vector **p** ∈ {0, 1}^*m*^ (or **p**_1_ ∈ {−1, 1}^*m*^) which indicates the underlying phenotype (disease status) for each genotype vector **g**_*j*_ also needs to be stored if the input VCF files are sent from the client in a random order.
ii. **Top** *l* **SNP candidate identification by sketching**. In this step the server keeps *d* independent sketches, each of size *w*, for approximate queries of 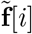, a linear proxy to the *χ*^2^ statistic 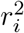 defined below. Specifically, let **g**_(i)_ denote the *i*-th column of the genotype matrix *G*, then 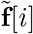 is given by the dot product of **g**_(i)_ and the phenotype vector **p**_1_ with *±*1 entries corrected by the top *y* eigenvectors 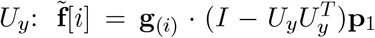 Let [*z*] = {1, *…, z*}. For updating the sketches the server maintains (i) *d* independent hash functions *h*_*k*_ (*k* ∈ [*d*]), where each hash function *h*_*k*_ : [*n*] → [*w*] maps a SNP *i* to one of the *w* buckets; and additionally (ii) *d* sign functions sign_1_, *…*, sign _d_ : [*n*] → {−1, 1}. Let *A*_*k*_ denote the array of size *w* of hash buckets that correspond to hash function *h*_*k*_. Initially all entries in the sketches *A*_1_, *…, A*_*d*_ are set to 0. To update the sketches, the server first precomputes a corrected phenotype vector with *±*1 entries 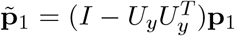, and then processes each encrypted VCF **g**_*j*_ and adds 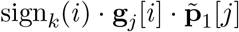 to *A*_*k*_[*h*_*k*_(*i*)], for each SNP *i* and hash function *k* ∈ [*d*]. Once all incoming data has been processed, the client sends the entire list of *n* SNP IDs to the server. For each SNP ID *i*, the server queries the *d* sketches and computes the median 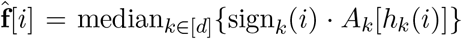. The top *l* SNP IDs according to the absolute value of 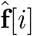 are kept for the next step. Ideally, these *l* SNPs include all top *k* SNPs according to the actual 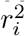 defined below.
iii. **Top** *k* **SNP identification among the** *l* **candidates** In this step we rank the *l* SNPs according to the Armitage trend *χ*^2^ statistic and return the *k* largest SNPs. The test statistic 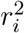 is defined for each SNP *i* as

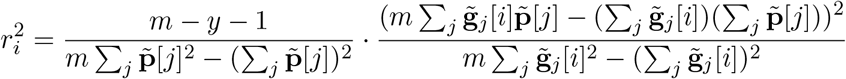

where 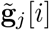 denotes the entries in a population stratification corrected genotype matrix 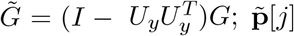 denotes the entries in a population stratification corrected phenotype vector 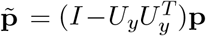. Rather than explicitly keeping the corrected genotype matrix 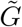 which requires *O*(*ml*) memory, SkSES implemented an algorithm which only maintains in the enclave *y* + 3 floating point numbers for each SNP *i* with a hash table (so the total memory required is *O*(*l · y*)), and these numbers are updated while processing the next individual **g**_*j*_. Once all *m* individuals are processed the Armitage trend *χ*^2^ statistic *r*_*i*_ for each SNP *i* can be computed exactly, and finally the top *k* SNPs according to *r*_*i*_ are returned to the client. We omitted the algorithmic details of how SkSES updates these numbers with **g**_*j*_ as this step is not modified with “learning-augmented” sketches described in the following section.

### 2.2 Our learning-augmented sketching approach for privacy preserving GWAS

We offer two major improvements over the SkSES method for (i) the PCA step and (ii) the top *l* SNP candidate identification step, by newly using learning-augmented sketches. To achieve this, we introduced a new pre-computing step, for which we assume that there is a small number of publicly available VCF files to be used as a training set; in case such a training set is not publicly available, a “small” proportion of the *m* VCF files in the input is sampled (with a default of *m/*10) and used as a training set. From the training set, the pre-computing step “learns” two sets of potentially significant SNPs in distinct phases: in the first phase, it computes *S*_1_ for assisting with SkSES’ step (i) and in the second phase, it computes *S*_2_, for assisting with SkSES step (ii). Note that both phases of the pre-computing step can be executed locally and outside a secure enclave. For example, when multiple computation parties are involved in a collaborative GWAS study, the sets *S*_1_ and *S*_2_ can be pre-computed in advance and locally by one of the computation parties, using the training set. Next, we modified steps (i) and (ii) of the SkSES’ secure GWAS protocol so that the sets *S*_1_ and *S*_2_ of potentially significant SNPs are sent to the server, along with all input data (i.e. preprocessed and compressed VCF files), for PCA and top *l* SNP candidate identification by sketching.

In the remainder of the paper we denote by *m*_0_ be the number of the VCF files in the training set and by *n*_0_ the total number of SNP records available in each one of the VCF files in the training set.

i. **Improved PCA with learned SNPs** The first phase of our new pre-computing step identifies the set of most significant SNPs *S*_1_ on the test set, i.e., genotype matrix 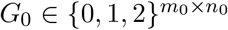. For that, we follow the same normalization process as SkSES to compute a mean-centered and standard deviation corrected genotype matrix *X*_0_ from *G*_0_. Then we compute the first eigenvector 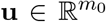 of the covariance matrix 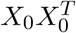 exactly. We pick the top *n′* SNPs as *S*_1_, according to their cosine similarities betwen each column vector in *G*_0_ and the top eigenvector **u** (recall that each SNP corresponds to a column vector in *G*_0_). Following the pre-computing step, the set *S*_1_ of *n′* SNP IDs is encrypted and sent to the server. Our newly modified step (i) of SkSES, now computes the top eigenvectors from the entire *m* samples within the secure enclave. For that, all participating parties send their encrypted VCF files to the server, which filters out all SNPs which are not present in *S*_1_, and performs SVD on the resulting genotype (sub)matrix *G* of dimension *m × n′*. Mean centering and normalization by row standard deviations before SVD (to compute *X* from *G*) are now done explicitly and in place. We note that *n′* by itself is not sensitive (in fact it is part of the GWAS protocol), and as such, we do not attempt to hide the message length of the encrypted SNP IDs sending to the server.
ii. **Improved top** *l* **SNP candidate identification with unique buckets**. In the second phase of our new pre-computing step, we identify the set of SNPs *S*_2_ by simply picking up *n″* SNPs with largest 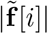 on the training set of *m*_0_ individuals among the *n*_0_ SNPs. *S*_2_ is then sent to the server just like *S*_1_ as described above. Our newly modified step (ii) of SkSES works as follows. First, the size of *S*_2_, *n″* is determined by the desired size of sketches to be maintained in the enclave memory, namely the product of *w*, the number of buckets in each sketch and *d*, the number of independent sketches: we split the *w · d* buckets originally assigned for sketches into two parts. The first *α · wd* buckets are allocated to maintain the SNP IDs as well as 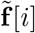 for each SNP *i* (this time, computed from the entire set of *m* individuals), through a hash table so that 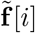 can be queried exactly. As such *n″* is given by 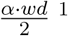. We refer to these buckets as “unique buckets” in the following and denote them as *B* [*i*] (*i* ∈ [*n″*]). The remaining (1 − *α*) *· wd* buckets are assigned to the sketches *A*_1_, *…, A*_*d*_, each of size reduced to (1 − *α*) *· w*. By default we set *α* = 0.5, meaning half of the buckets originally used for maintaining sketches in the enclave memory are assigned as unique buckets *B*[*i*]. To process an encrypted VCF file **g**_*j*_ that includes a SNP *i*, the server first checks whether the SNP ID *i* is maintained in the unique buckets. If yes, then we add 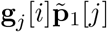 to the unique bucket *B*[*i*], otherwise update the sketch entries 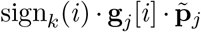 to *A*_*k*_[*h*_*k*_(*i*)] for each *k* ∈ [*d*]. After all incoming VCF files has been processed, the client again sends the entire list of *n* SNP IDs to the server. The server modified its querying process for SNP *i* by also first checking whether *i* lies in the unique buckets. If yes, then the query returns the value currently stored in *B*[*i*] as the exact 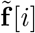; otherwise sketch entries are queried and an approximate 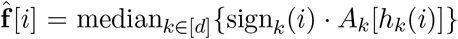 is returned. Still the *l* largest SNPs according to the absolute value of 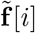 (or 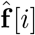) are stored as candidates for the top *k* SNP identification in the next step.

#### Comparison with existing work on learning augmented algorithms

There are a number of recent works in using machine learning to augment the performance of classical algorithms [31, 32, 33, 34, 35, 36]. Unlike the most relevant *Learning-enhanced Frequency Estimation Algorithms* [32], where the IP addresses carry meaningful features, we cannot extract meaningful features from the SNP IDs (rsid) alone. We do not have enough memory to store auxiliary data inside the enclave, and querying an external database would leak information. So we resort to memorization rather than generalization, i.e. we learn explicit sets of SNPs rather than an oracle to classify the SNPs.

## 3 Results

### 3.1 Dataset

We evaluated our approach on a dataset from the UK Biobank consisting of 7,550 GVCF files, with 3,775 from type 2 diabetes patients^2^ (cases), and the other 3,775 from non-patients (controls). These GVCF files include in total 9.54 *×* 10^5^ distinct SNPs, and we performed imputation on each individual with the 1000 Genomes Phase 3 [37] reference panel which includes 7.82 *×* 10^7^ distinct SNPs in the human genome. We sampled random subsets of 100, 200, 400, 800, 1600 and 3200 imputed GVCF files to use them as training sets for our learning augmented approach, and then applied it to test sets composed of, again randomly sampled 1,000, 2,000, and 4,000 imputed GVCF files (the training and test sets did not overlap and they were both composed 50% cases and 50% controls) to evaluate the performance of our approach.

Note that each of the original 7,550 GVCF files from the UK Biobank contains ∼ 80k variant records. Before using them as inputs to our approach we processed each GVCF file by first phasing its variants using SHAPEIT4 [38] and then imputed the missing allele specific variants using Minimac [39]. After imputation, each resulting VCF ended up containing genotype information of 7.82 *×* 10^7^ variants as mentioned above. Interestingly only 2.35 *×* 10^7^ of the variants were present in at least one of the 7,550 individuals, so the remainder of the variants were removed from the VCFs. Therefore, all training and test sets we used were comprised of 2.35 *×* 10^7^ (*n* ≤ 2.35 *×* 10^7^) variant records. Please see the supplementary materials for further details about how we prepared our datasets.

### 3.2 Tested methods and parameters

We tested our learning-augmented sketching method against the **SkSES** baseline on the datasets described below. We report the results from our main **fully-learned** approach, where we utilize both sets of learned SNPs *S*_1_ (for PCA) and *S*_2_ (for unique buckets), as well as three different approaches that each uses only part of the full method to study their individual effect: the **learned PCA** approach where we only utilize *S*_1_, the **learned unique** approach where we only utilize *S*_2_, and the **no sketch** approach, where both *S*_1_ and *S*_2_ are used but the sketch is completely removed in favor of the unique buckets.

For all four learning-augmented sketching methods, we ensured that the training set was evenly split between cases and controls just like the full dataset it was sampled from. The original SkSES method does not use a training set. We set the size of the set *S*_1_ of learned SNPs (for PCA) to be *n′* = 4,096, consistent with the sketch matrix dimension used in the original SkSES. We used two principal components for population stratification correction (*y* = 2), the default setting of SkSES. We set the number of candidates *l* to be 2^20^. The size of training set *m*_0_, the size of sketches *w*, and the number of unique buckets *n″* vary between different experiment settings.

### 3.3 Memory and time consumption

We first compared the runtime and memory consumption of the learning-augmented sketching approach (fully-learned) with that of SkSES. We ensured that the fully-learned method uses the same amount of memory as the original SkSES algorithm. Numeric values are stored as 32-bit floating numbers, occupying 4B of memory each. The memory requirements of the three steps described in Section 2.1.2 are as follows. (i) PCA requires *m · n′ ·* 4B=62.5MB memory. (ii) Sketching memory requirements depend on other parameters. For SkSES, we chose the parameters *d* = 8 and *w* = 2^21^ for the sketching arrays, occupying 64MB memory in total. For our learning-augmented sketching method, we used sketching arrays with parameters *d* = 8 and *w* = 2^20^ (32MB) together with *n″* = 2^22^ unique buckets (32MB). (iii) Computing the *χ*^2^ values accurately for the *l* candidates requires *l*(3 + *y*) *·* 4B=20MB.

Note that, while transitioning from step (ii) to (iii), and before the sketching arrays and unique buckets are cleared, each approach (including SkSES) requires and additional 2*l ·*4B=8MB memory. Therefore, during all our tests the memory used by each method at any point of execution was at most 72MB, ignoring a small overhead. This means that the parameters *d, w, n″*, and *l* could be further increased, but how we should increase them for optimal performance while remaining within 128MB or a higher upper bound would be the subject of future work.

In our tests, our learning augmented method takes 1,978 seconds to complete step (i), 4,419 seconds to complete step (ii) (which are significantly faster than the original SkSES algorithm), and 1,323 seconds to complete step (iii) (see Table 1). These numbers do not include the the time for training and preprocessing (including compression and encryption) the VCF files, which can be done locally by each participating party individually with low cost. The tests were run on a Linux server equipped with an SGX-capable Intel Xeon E3-1280 v5 processor with 4 cores at 3.70 GHz.

**Table 1:** Running time performance of learning-augmented sketching method (fully-learned) against SkSES, with identical enclave memory allocated to core data structures. Mean running time *±* standard deviation is reported from the results of five runs. In step (i) popstrat correction using PCA, we set *n′* = 4096 for both methods. In Step (ii) top *l* SNP candidate identification by sketching, we set *d* = 8 and *w* = 2^21^ for the sketch in the SkSES method; *d* = 8 and *w* = 2^20^ for the sketch together with *n″* = 2^22^ unique buckets for learning-augmented sketching method, so that both methods use 64MB memory. In step (iii) top-*k* SNP identification among *l* we set *l* = 2^20^ for both methods. In our implementation of the fully-learned method, we replaced the Robin Hood hash table (used in step (iii)) in the original SkSES by a plain array to fix a bug, which results in a slowdown. If the fix is directly applied to the original SkSES, step (iii) slows down to 1255*±*13s.

| Step/Method | SkSES | fully-learned |
| --- | --- | --- |
| (i) Popstrat correction using PCA | 3846± 16s | 1610± 10s |
| (ii) Top-$l$ SNP candidate identification by sketching | 8767±100s | 4294±222s |
| (iii) Top-$k$ SNP identification among $l$ candidates | 32± 0s | 1240± 11s |

We additionally tested the running time performance with varying sketch width *w* and depth *d*, and concluded that the time consumed by step (ii) appears to be determined mostly by *w*; in fact starting from *w* = 2^17^ the time roughly doubles as *w* doubles, even though the amount of read/writes remain the same. Decreasing *w* by increasing *d* or introducing unique buckets is therefore beneficial for reducing running times.

### 3.4 Accuracy of sketches

We performed three sets of experiments to assess our learning augmented sketching approach. In the first set of experiments we assessed the impact of the training set size, in comparison to the test set size. In the second set of experiments we assessed the impact of the size of the sketches and the number of unique buckets. In the third and final set of experiments we assessed how our learning augmented sketching approach compares against the baseline approaches and how individual components of the full approach contribute to its performance for various test set sizes.

The metric we used to evaluate performance of the algorithms is the true positive rate, defined to be the percentage of the top *k* SNPs identified that are actually among the top *k*, according to the Cochran–Armitage trend *χ*^2^ statistic. We report the percentage for both *k* = 100 (with Cochran–Armitage *r*^2^ = 19.61/20.86/23.43 from 1000/2000/4000 individuals) and *k* = 1,000 (with Cochran–Armitage *r*^2^ = 15.15/14.77/17.91 from 1000/2000/4000 individuals).

#### 3.4.1 Impact of training set size

In the first set of experiments we assessed the impact of the training set size on accuracy, given the test set size. We used the following settings in these experiments (which, as per Table 3, provide the best results): the number of unique buckets are 2^22^, the sketch depth is 8 and the sketch width is 2^20^. Note that we used training sets disjoint from the tests sets.^3^ As demonstrated in Table 2, the impact of training set size on the accuracy is minimal, as long as the training set size is at least 200. In fact, if the whole test set is used as the training set, we do not get much (or any) improvement: for the 1,000-individual test set we get 1.000/0.899; for the 2,000-individual test set we get 0.990/0.824; for the 4,000-individual test set we get 0.940/0.788.

**Table 2:**
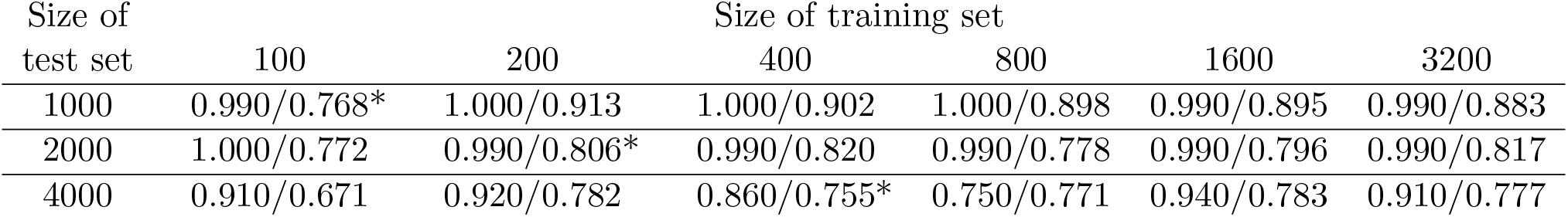
Accuracy of the fully-learned method (with 2^22^ unique buckets and a sketch of depth 8 and width 2^20^) as a function of training set size. Note that the training sets were sampled from 3550 individuals disjoint from test sets with 1000*/*2000*/*4000 individuals. With each training set (column) and each test set (row) size, we give the accuracy values for both the top-100 (left) and top-1000 (right) SNPs. The three entries marked with * are the configurations used in the (Figures in the) remainder of the paper.

**Table 3:** Accuracy of the fully-learned approach under various configurations for the sketch width (rows) and the number of unique buckets (columns) on a test set with 4000 individuals trained on an independent, randomly selected set with 400 individuals (sketch depth: 8). For each number of unique buckets (column) and each sketch width (row), we give the accuracy values for both the top-100 (left) and top-1000 (right) SNPs. The results in the rightmost column are those with no (learned) unique buckets but with learned PCA. The same amount of memory is used for the unique buckets as the sketch for entries along the main diagonal (except for the rightmost column). The entry marked with * is the configuration used in the (Figures of the) remainder of the paper; it is the first diagonal entry that fits within 128MB SGX enclave memory.

| Sketch width<br>($w$) | Number of unique buckets ($n''$) | | | | |
| --- | --- | --- | --- | --- | --- |
| | $2^{23}$ | $2^{22}$ | $2^{21}$ | $2^{20}$ | 0 |
| $2^{21}$ | 0.940/0.791 | 0.950/0.784 | 0.790/0.753 | 0.730/0.697 | 0.590/0.653 |
| $2^{20}$ | 0.940/0.788 | 0.900/0.770* | 0.650/0.672 | 0.600/0.635 | 0.470/0.555 |
| $2^{19}$ | 0.940/0.785 | 0.570/0.668 | 0.540/0.611 | 0.500/0.543 | 0.400/0.440 |
| $2^{18}$ | 0.920/0.769 | 0.490/0.580 | 0.440/0.465 | 0.350/0.440 | 0.240/0.318 |
| $2^{17}$ | 0.440/0.602 | 0.410/0.484 | 0.260/0.349 | 0.180/0.288 | 0.120/0.205 |

**Table 4:**
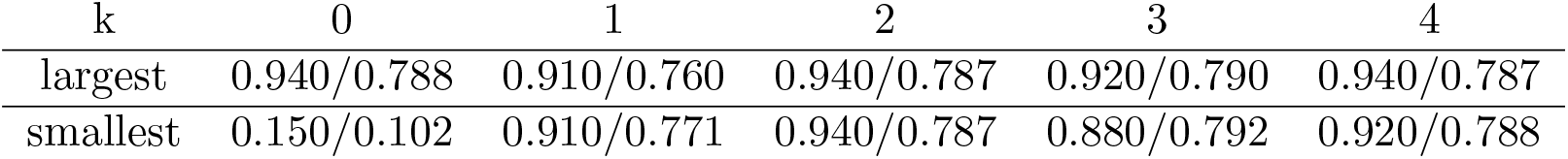
Accuracy of the fully-learned approach using a non-representative training set. For each *k*, the training set consists of the individuals *j* in the cohort, with the largest (middle row) or smallest (bottom row) 400 entries (**u**_*k*_)_*j*_ in the *k*^*th*^ eigenvector. For a given *k*, the accuracy of the fully-learned approach for the top-100/top-1000 SNPs are given.

#### 3.4.2 Impact of sketch size and unique bucket count

In the second set of experiments we assessed the impact of the sketch size vs the number of unique buckets on accuracy (training set size: 400, test set size: 4,000). As demonstrated in Table 3, for a given number of unique buckets, there is a sharp drop in accuracy as the sketch width decreases, probably because when the sketch is small, the values accumulated in the sketch and hence the queried values are systematically larger (in absolute value) than the values in the unique buckets, leading to few or none of the SNPs in unique buckets being selected as the top *l* candidates. We leave a more comprehensive study of methods to balance the sketch size and the number of unique buckets to future work.

#### 3.4.3 Comparison with baseline and ablation study

In the next set of experiments, we compared the accuracies of the four learning-augmented approaches with the SkSES baseline. Results from five runs of each method are shown as box plots in Figures 1, 2 and and 3, except for the “no sketch” approach, which is only run once because no randomness is involved. All other methods use random hash functions for the sketch, and the “learned unique” and SkSES methods also use a different random seed for each run to generate the sketch matrix *K* for PCA. The training sets stay the same across five runs.

**Figure 1.**
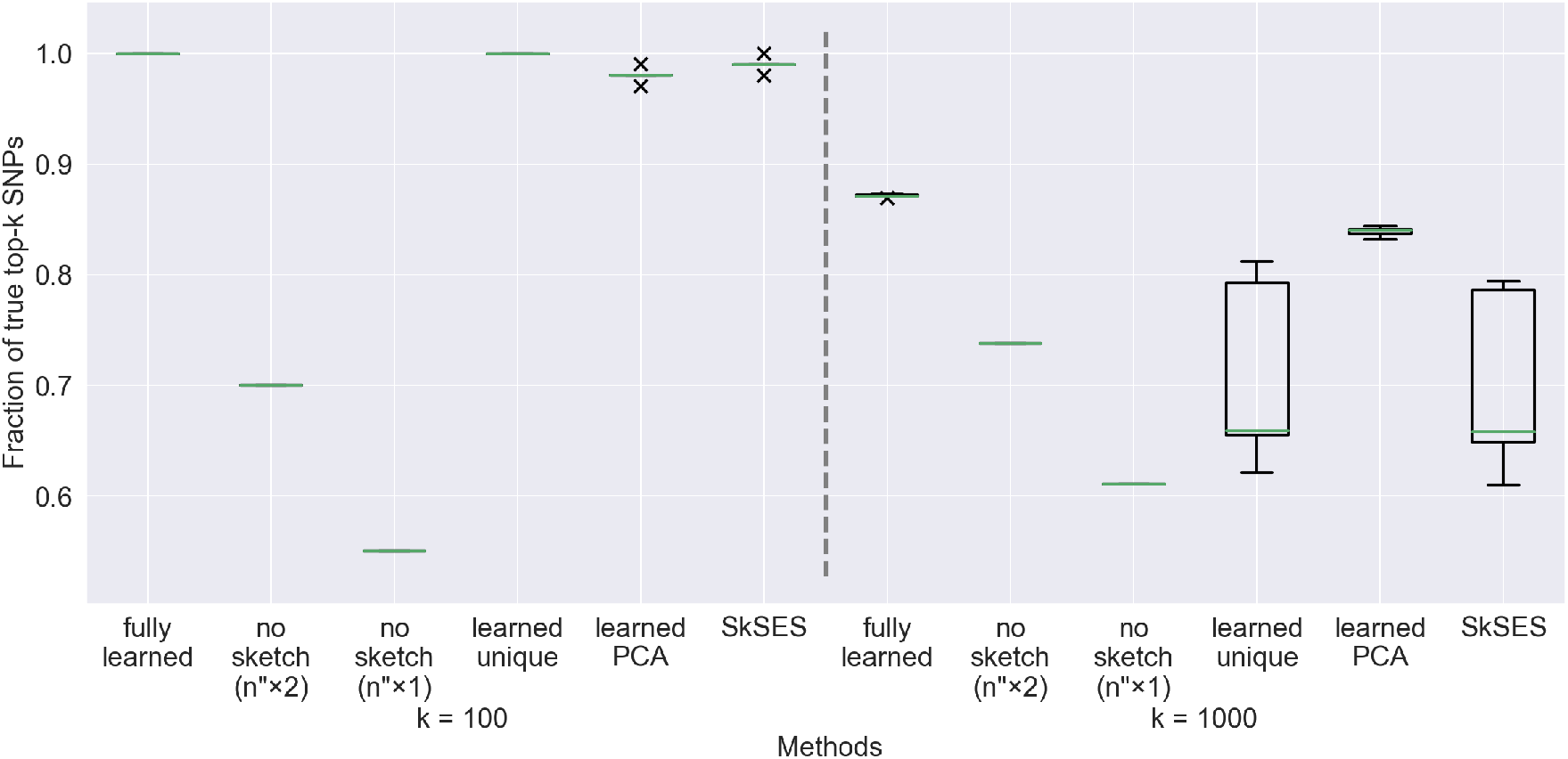
Accuracy comparison of the five approaches on a dataset of 1,000 individuals, using a training set of 100 individuals. The percentages of the actual top 100 and top 1,000 SNPs identified are shown. Both the “learned unique” and SkSES methods use a randomly signed sketch matrix *K* generated by a random seed, and we observe that some random seeds lead to exceptionally good results (for example the outliers near 0.8 in with *k* =1,000 for both methods); these seeds appear roughly 20% of the time when randomly selected.

**Figure 2.**
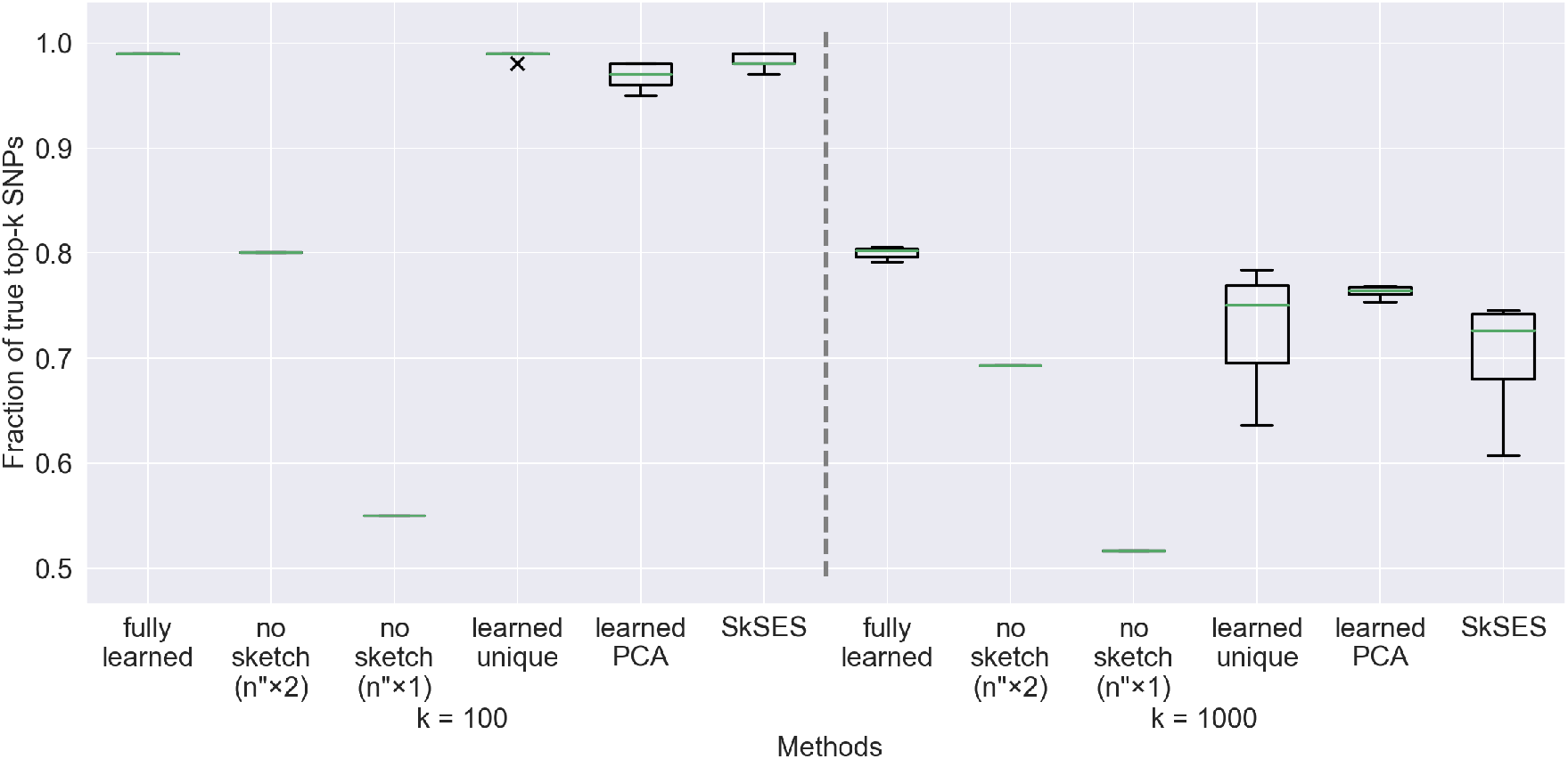
Accuracy results on a dataset of 2,000 individuals using a training set of 200 individuals. The large variations in accuracy of the “learned unique” and SkSES approaches are due to the use of a random seed to generate the sketch matrix *K*.

**Figure 3.**
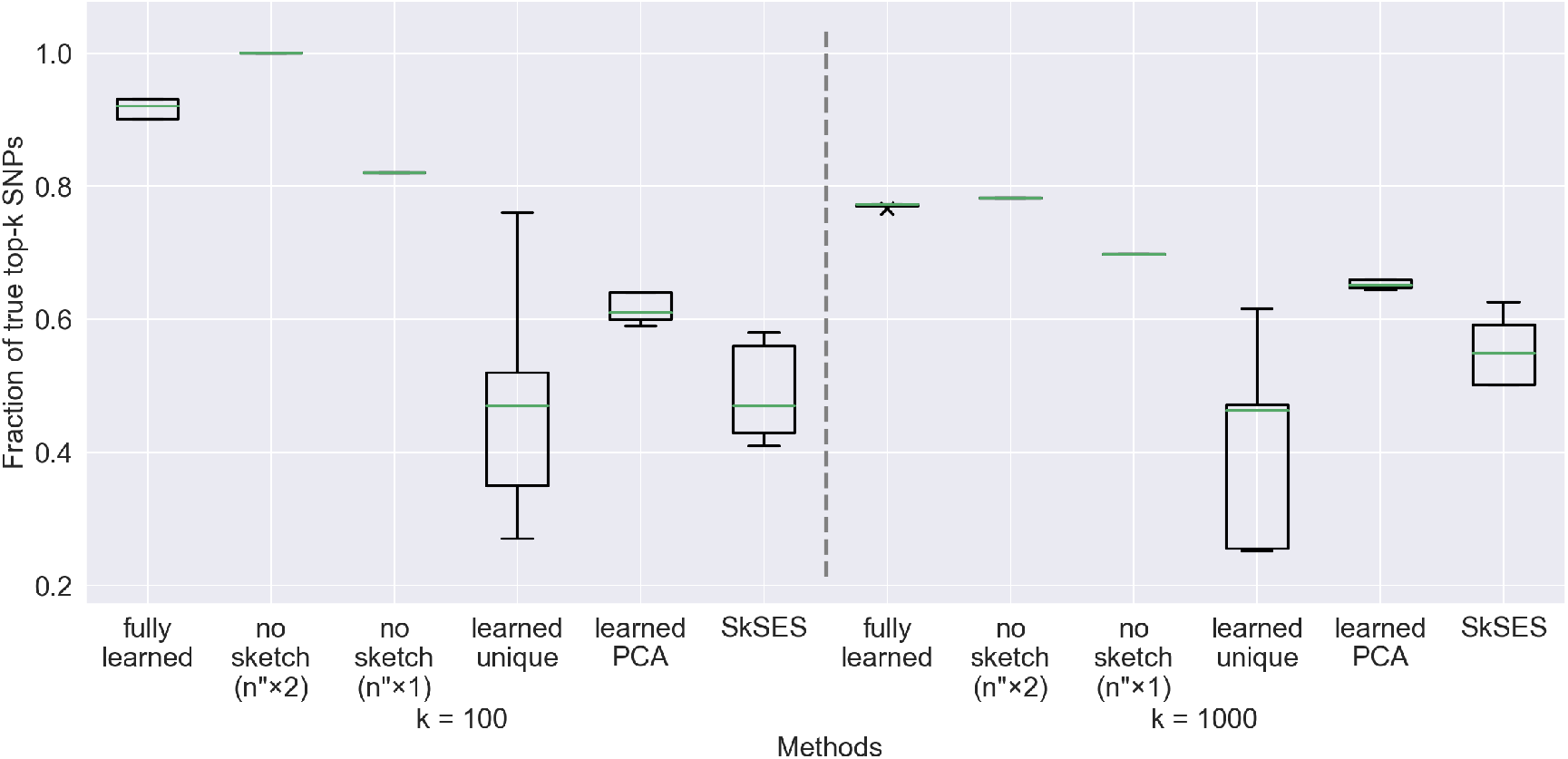
Accuracy results on a dataset of 4,000 individuals using a training set of 400 individuals. The large variations in accuracy of the “learned unique” and SkSES approaches are due to the use of a random seed to generate the sketch matrix *K*. Notice that the “fully learned” method is not superior to “no sketch” under memory parity (*n″ ×* 2) here (while it is superior on the 1,000- and 2,000-individual test sets), but the usefulness of sketching can still be seen when compared with “no sketch” (*n″ ×* 1).

For all four learning-augmented methods, the training set size is fixed to be 10% of the test set size, which can be compared against entries marked with * in Table 2. We ensure all methods use the same amount of memory in step (ii) in order to compare them fairly: the sketch depth and width for the main approach (fully-learned) are respectively fixed at 8 and 2^20^, and the unique bucket count is fixed at 2^22^, which is marked with * in Table 3. For the other approaches, if sketching is not used (“no sketch”) then the unique bucket count *n″* is doubled (fixed at 2^23^), while if unique buckets are not used (“learned PCA” and SkSES) then the sketch width is doubled (fixed at 2^21^). Since we also report the performance of the “no sketch” approach without doubling the unique bucket count in the same figures - to assess the accuracy gains that can be attributed to sketching, we distinguish the two setups by their labels: “no sketch (*n″ ×* 2)” v.s. “no sketch (*n″ ×* 1) respectively.

Under the same memory constraint, our fully-learned approach scored a 40% absolute improvement (from ∼ 50% to *>* 90%) of the percentage on the 4,000-individual test set when *k* = 100, and a 20% absolute improvement of the percentage when *k* = 1,000, compared to the original SkSES algorithm (our baseline); see Figure 3. The improvement on the 2,000- and 1,000-individual test sets are less pronounced but still clearly visible, as shown in Figure 1 and Figure 2.

It is crucial to utilize the small set *S*_1_ of significant SNPs in the PCA step to create the sketch matrix *K* for PCA, which leads to major improvement and stabilization of performance. If we used a randomly signed sketch matrix as in the original SkSES (the “learned unique” approach), the performance is very unstable: for about 10% of such randomly chosen matrices, we get performance matching the performance of “fully learned” approach, but most of the time we get very poor results. It is still an open question why using a subset of SNPs that is order-of-magnitudes smaller than *n* yields more stable eigenvector results than using the whole 23 million SNPs in a sketched form, and we leave this to future work.

### 3.5 Using a non-representative training set

To show robustness of our method, we tested its performance using training sets that are not representative of the whole test set. Specifically, we selected 400-individual subsets “of different ethnicities” from the 4000-individual test set in the following way: for *k* = 0, 1, 2, 3, 4, we take the *k*-th eigenvector **u**_*k*_ (where *k* = 0 corresponds to the top eigenvalue) of the covariance matrix *X′X′*^*T*^ of the normalized genotype matrix *X′*, and pick the 400 individuals *j* with (i) the largest, and then (ii) the smallest (**u**_*k*_)_*j*_, and use them to form the training set.^4^ As shown in the table below, the results are similar to those using a disjoint (*§*3.4.1) or sampled (*§*3.4.3) training set. For the 400 individuals with the smallest (**u**_*k*_)_*j*_, the result is quite poor for *k* = 0 (corresponding to the top eigenvector, presumably representing the most common ethnic contribution to this cohort), but remains good for *k* = 1, 2, 3, 4.^5^

## 4 Acknowledgements

We thank the Department of Computer Science at Indiana University, especially Haixu Tang and Xiaofeng Wang, for providing remote access to SGX-enabled machines on which we run the experiments. We also thank Hongbo Chen and Rob Henderson for providing technical assistance.

This research was supported in part by the Intramural Research Program of the Center for Cancer Research, National Cancer Institute, NIH.

## 6. Supplementary materials: dataset preparation

**Original data**. The SNPs in UKB data were called from whole exome sequencing and are provided in compressed GVCF format with suffix `gvcf.gz`, where each line contains an additional `<NON_REF>` indicator in addition to the `ALT` allele (or only `<NON_REF>` in the `ALT` field). These `<NON_REF>` indicate “blocks” of `0/0` genotypes: their `INFO` fields have the form `END=`some number, which indicate there are no variants called within the range from `POS` to `END`.^6^.

The SNPs in the GVCF files were called using the reference genome GRCh38. Each GVCF contains about 2 million SNP records, but most of them are `<NON_REF>` only (indicating `0/0` blocks without variants), and there are only about 80k explicit variant records. Due to whole exome sequencing, each of the VCF files covers only 3 ∼ 5% of the whole genome length. Most heterozygous genotype fields (`GT`) are not phased, shown as `0/1` instead of `0|1` or `1|0`. Some genotypes are phased (shown as `PGT`) relative to a nearby variant (`PID`), probably called by GATK when the variants simultaneously appear in some reads.

### Data augmentation via phasing and imputation

We ran SHAPEIT4 [38] and Minimac4 to impute the set of 9.54 *×* 10^5^ whole exome SNPs to whole genome (including introns), so our learning-augmented sketching method can run on a dataset of larger scale where the memory saving is of more significance. Specifically we first phased the UKB dataset using SHAPEIT4 and then imputed the resulting files using Minimac4, with the following custom commands.

Minimac4 can read the `gvcf.gz` files directly, but SHAPEIT4 requires the input to be compressed using bgzip, and indexed using either Tabix (`.tbi`) or bcftools (`.csi`), so we need to decompress, recompress with bgzip, and index. None of the tools have native support for GVCF: none can recognize the `<NON_REF>` indicators, and they must be removed (or replaced with a dot when `<NON_REF>` is the only one) during this preprocessing step. The `POS`-`END` blocks were not recognized either: the variants in the reference panel that lie within the blocks do not contribute to the *overlap* between input and reference variants as reported by any of the tools. If `<NON_REF>` were not removed, 0 overlap will be reported. For Chromosome 1, there are roughly 6k SNPs overlapping with the GRCh38 reference panel; but if the wrong reference panel GRCh37 were used for imputation/phasing, this number will drop to *<* 20. We note that these blocks of `0/0`s need to be converted to explicit lines of variant records (that exist in the reference panel) via a custom script, since we were unable to find an existing and publicly available tool that conveniently does this. After this conversion, the overlap raised from 6,212 to 237,082 for Chromosome 1 for the first individual among the 3775 cases, a 37-fold increase. Without this conversion, Minimac4 imputation might output nonzero values for variants lying in the `0/0` blocks. For example, 659 variants were output for the first individual within the `0/0` blocks.

The default 1000 Genome reference panel provided by Minimac at https://genome.sph.umich.edu/wiki/Minimac3#Reference_Panels_for_Download cannot be used because it uses reference GRCh37. Instead, we downloaded the GRCh38 reference panel (also from 1000 Genome Phase 3) from http://hgdownload.soe.ucsc.edu/gbdb/hg38/1000Genomes/, which are bgzipped VCF files indexed with Tabix. This reference panel can be directly used by SHAPEIT4, but for Minimac4, we need to convert it to m3vcf with Minimac3, using the command

~~~
minimac3 --refHaps ALL.chr1.shapeit2_integrated_snvindels_v2a_27022019.GRCh38.phased
.vcf.gz --processReference --prefix chr1
~~~

for chromosome 1, for example.

Both SHAPEIT4 and Minimac4 process one chromosome at a time, so it is necessary to use VCFtools [40] to extract one chromosome at a time. SHAPEIT4 requires an additional genetic map file, which can be downloaded from https://github.com/odelaneau/shapeit4/tree/master/ maps, b38. Minimac4 requires input/target VCF to be pre-phased and is unable to phase by itself; SHAPEIT4 can phase and impute but is designed specifically for high-accuracy phasing. Minimac4 supports multi-allelic sites in reference panel, and keeps the original variant records. We ignored all pre-phased sites marked by `PGT` (local/partial phasing) in the original GVCFs when running SHAPEIT4.

### Example commands

As an example, given an unphased VCF file `case_0_chr1.vcf`, we first use our custom script `block_conv.py` to convert 0/0 blocks to explicit variant records. Then we use BGZIP to compress it and BCFtools to index it, and run SHAPEIT4 with the following commands:

~~~
bgzip -c case_0_chr1.vcf > case_0_chr1.vcf.gz
bcftools index case_0_chr1.vcf.gz
shapeit --input cases/case_0_chr1.vcf.gz \
--map shapeitmap/b38/chr1.b38.gmap.gz --region 1 --reference \
snv-indel-ref/ALL.chr1.shapeit2_integrated_snvindels_v2a_27022019.GRCh38.phased.vcf.gz \
--output case_0_chr1_phased.vcf.gz
~~~

SHAPEIT4 reports an overlap L=237,082, performs phasing and outputs 230,880 records with genotype `0|0`, 2,006 with `0|1`, 1,779 with `1|0`, 2,417 with `1|1`, for a total of 237,082. (If 0/0 blocks were left unconverted, it reports a mere overlap of L=6,212 and outputs 10 records with `0|0`, 2,011 with `0|1`, 1,774 with `1|0`, and 2,417 with `1|1`.)

Once phasing is done, we run Minimac4 with the following command:

~~~
minimac4 --refHaps snv-indel-ref/chr1.m3vcf.gz --haps case_0_chr1_phased.vcf.gz \
--prefix minimac_imputed --ChunkLengthMb 300 --rsid --allTypedSites
~~~

In the resulting imputed VCF file, Minimac outputs 5,957,753 records with genotype `0|0`, 54,723 with `0|1`, 51,132 with `1|0`, and 128,225 with `1|1`, for a total of 6,191,833.

## Footnotes

1 we ignore the cases where *n″* ≥ *n*_0_ as the assumption is the total number of SNPs is significantly larger than the size of the sketch, and the *m*_0_ samples should include sufficiently large number of SNPs to be tested

2 the entire set of type 2 diabetes patients with relevant information in the UK Biobank

3 There is some minimal impact if the training sets were sampled from the test sets: a comparison of the three cells marked with * with the corresponding Figures reveal that only the 1,000 individual case has an increase in accuracy when the training set is indeed sampled from the test set.

4 Since **u**_*k*_ is an eigenvector of the real symmetric matrix *XX*^*T*^, we have *XX*^*T*^ **u**_*k*_ = *λ*_*k*_**u**_*k*_ for some nonnegative number *λ*_*k*_, so (**u**_*k*_)_*j*_ is proportional to the dot product of *j*th individual’s correlation vector with **u**_*k*_.

5 Since the dataset is from the UK Biobank, it is possible that there is a dominant ethnicity in the cohort and the selection of a training subcohort with the lowest contribution from this dominant ethnicity significantly impacted these results, especially because the SNP set *S*_1_, which we use for performing PCA, consists of those SNPs with the highest cosine similarity with the top eigenvector.

6 cf. https://gatk.broadinstitute.org/hc/en-us/articles/360035531812-GVCF-Genomic-Variant-Call-Format, “The two types of GVCFs” and “Records”, and https://samtools.github.io/hts-specs/VCFv4.3.pdf, p.9

